# Assessment of dynamics of global DNA methylation during the cryopreservation process of *Pinus pinaster* embryogenic lines

**DOI:** 10.1101/2022.02.27.481657

**Authors:** Isabel Mendoza-Poudereux, María Cano, María Teresa Solís, Francisco Esteve-Díaz, Pilar S. Testillano, Juan Segura, Ester Sales, Isabel Arrillaga

## Abstract

Long-term in vitro maintenance of embryogenic lines of *Pinus* species has been associated with lower maturation capacity, because of this, cryopreservation protocols for the embryogenic lines are needed to maintain valuable genotypes. Since cryopreservation may induce epigenetic variations, we evaluate changes in DNA methylation levels through the course of the cryopreservation of maritime pine embryogenic lines, as compared to those lines maintained by repeated subcultures. Six maritime pine embryogenic lines were cryopreserved following a protocol that includes pre-treatments inducing osmotic stress in liquid media. The percentage of methylated cytosines (%5-mC) in total DNA was determined by using a colorimetric assay. Reactive oxygen species (ROS) accumulation in cell lines was also determined by quantifying dihydroethidium (DHE) fluorescence under a confocal laser-scanning microscope. In the first experiment, we found that global DNA methylation was significantly reduced during the cryopreservation protocol. Subsequently, we evaluated the methylation status of both cryopreserved and no cryopreserved lines (maintained by subcultures) and found differences among embryogenic lines but overall, cryopreservation did not alter %5-mC of the recovered lines while periodical subcultures increased methylation rates. In addition, maltose pretreatment did not increase significantly ROS production in embryogenic lines. Our results demonstrate that the genetic stability during cryopreservation highly depends on the embryogenic line studied, but the protocol allows maintaining methylation DNA rates in most of the recovered lines.

## Introduction

*Pinus pinaster* Aiton is a coniferous species native to the Southern Europe with a high economic and recreational value. It is considered a model species for the study of trees’ adaptation responses to drought and nutritional stresses (Arrillaga et al. 2014). In vitro somatic embryogenesis (SE) is the most efficient biotechnological approach for conifer clonal propagation, and then useful for research and breeding (Klimaszewska et al 2018, Lelu-Walter 2016). Nevertheless, maritime pine embryogenic lines lose maturation capability after a prolonged period in culture (Humánez et al. 2012). To date, the genetic stability of embryogenic lines from maritime pine has only been evaluated by SSR markers, which detected variation in this plant material under proliferation conditions (Marum et al. 2009). For the appropriate conservation of selected lines, cryopreservation in liquid nitrogen (LN) is therefore required. Cryopreservation may allow safekeeping of rare and valuable germplasm, providing also a potential backup of clonal lines while trees are tested in the field (Park 2002). Marum et al. (2004) described a protocol for cryopreservation of embryogenic lines from maritime pine routinely used in many laboratories, but genetic stability during the cryopreservation process has not been evaluated so far. In some cases, the *in vitro* culture procedures required in a cryopreservation protocol have been considered the main origin of the detected changes, including epigenetic variations (Ibáñez et al. 2019).

Genome stability in plants and mammalians is regulated by DNA methylation, a conserved epigenetic mark also involved in gene expression and gene imprinting; thereby, disruption of DNA methylation can lead to developmental abnormalities (Gallego-Bartolomé 2020, Zhang et al. 2018). DNA methylation refers to the addition of a methyl group to the cytosine bases of DNA to form 5-methylcytosine (Tirnaz and Batley 2019) and participates in shaping the structure of the genome, modulating gene expression by inhibiting proteins binding to DNA, and by changing the structure of the associated chromatin (Niederuth and Schmitz 2017). In plants, cytosine-methylation can occur in any context (CG, CHG and asymmetric CHH, where H is A, C or T) with CG being the most commonly methylated dinucleotide (Osabe et al. 2014). A correlation between methylation of specific cytosine nucleotides and the frequency of point mutations at these sites has been described (Kovalchuk 2016), and also epigenetic changes have been associated to genetic variations, or considered precursors of somaclonal variation (Ibáñez et al. 2019).

Many technologies have been developed over the past decade to measure DNA methylation. Some of them use proteins that selectively bind methylated cytosines, while in others DNA is digested with methylation-sensitive restriction enzymes before amplification (methylation-sensitive amplified polymorphism, MSAP). Next-generation sequencing-based methods are also used, after conversion of unmethylated cytosines to uracils using sodium bisulfite (whole-genome bisulfite sequencing, WGBS). The selection of the most appropriate assay depends mainly on the aim of the study (profiling whole genome methylation or to find differentially methylated regions or candidate genes), as well as on the quantity and purity of the available DNA (Kurdyukov and Bullock 2016).

During cryopreservation, cells may suffer many damages, such as osmotic dehydration, large ice puncture and oxidative damages from reactive oxygen species (ROS). Classical cryoprotectants are employed to reduce osmotic and mechanical damages, but fail to dispose ROS (Liu et al. 2021). Furthermore, cryoprotectants used to support cell resistance to cryoinjury can cause oxidative stress at high concentrations (Ren et al. 2015). Maintaining ROS homeostasis is a key factor for avoiding oxidative damage, but ROS also function as important signaling molecules that regulate normal plant growth and responses to stress, and interplay with epigenetic modifiers and hormones, although the regulatory mechanism of that remains unknown (Huang et al. 2019).

With the aim of to know whether cryopreservation induce epigenetic changes in *Pinus pinaster,* we evaluated global DNA methylation and ROS production in embryogenic lines of the species during the cryopreservation process and after thawing and further proliferation.

## Materials and methods

### Plant material

Six maritime pine (*Pinus pinaster* Aiton) embryogenic lines (7P9a, 58P1a, 58P5a, B5, B14, and B50) were used in the analyses. Embryogenic lines were generated from megagametophytes collected from 5 open-pollinated mother trees and maintained on mLV medium (Lelu-Walter et al. 2006) with 2 week-subcultures, as described in Cano et al. (2019). Three-month-old embryogenic lines were cryopreserved following a protocol based in Marum et al. (2004), modified as described in Cano et al. (2019) and depicted in Figure 1. Aliquots were sampled at each step of the process to study changes in global DNA methylation level. Briefly, 3 g of actively growing embryogenic cell masses (10 days after the last subculture) were disaggregated in 10.8 mL of mLV medium in the dark for 24h at 90 rpm (sample T24); then, maltose was added until a 0.2 M concentration was reached. After 24 h, an aliquot was taken (sample T48) and maltose concentration was increased up to 0.4 M. On the third day, a new aliquot was taken (T72) and the flasks were transferred for 15 min to ice, and 12.6 mL of PSD solution (mLV medium supplemented with 10% Sucrose, 10% PEG-4000 and 10% DMSO) were added. Aliquots (1.8 mL) were distributed in cryovials, placed in a Nalgene® Cryo 1°C “Mr. Frosty” Freezing Container (filled with isopropyl alcohol and previously incubated at −80°C for 24 hours) and kept in an ultrafreezer at −80°C.

**Figure 1.**
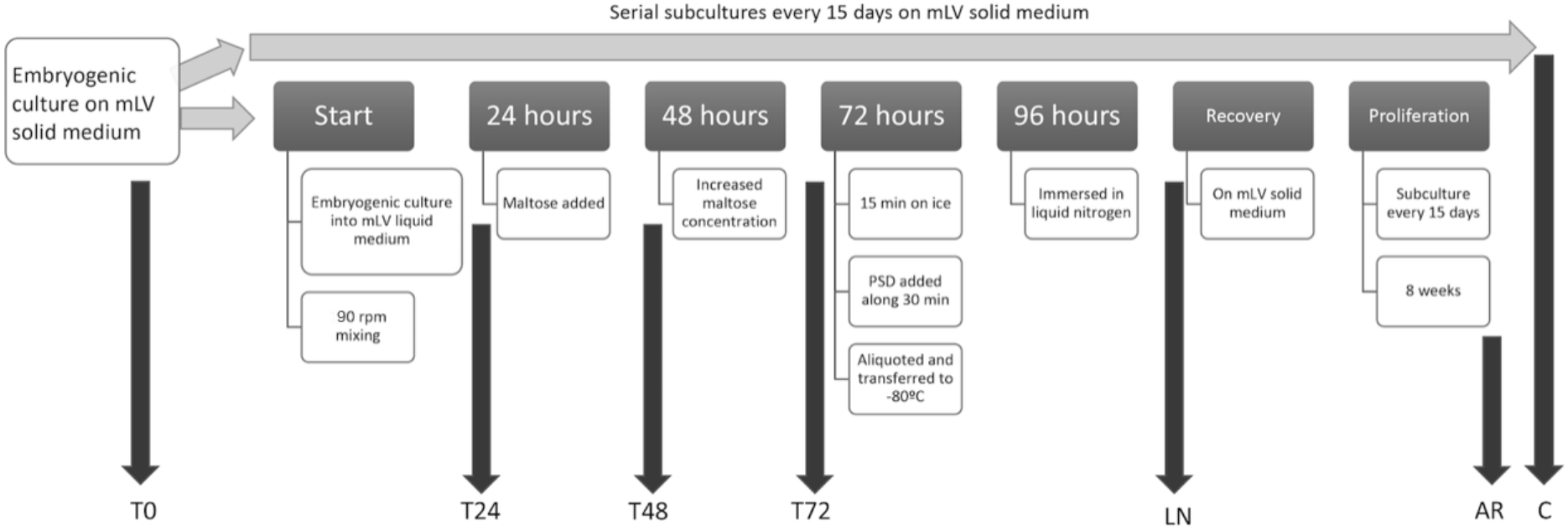
Cryopreservation procedure and sampling points (arrows). T0, before cryopreservation; T24, after 24h in liquid medium before maltose addition; T48, after 24h in 0.2 M maltose; T72, after increasing maltose up to 0.4 M; LN, after being submerged in liquid nitrogen; AR, after 8-weeks recovery; C, no cryopreserved lines after 8 weeks.

Cryovials were taken from the −80 °C ultrafreezer after 24 h and dipped into liquid nitrogen (LN, −196 °C) for their storage. After 24 hours in LN, cryovials were recovered (sample LN), shortly placed into a bath at 40 °C and then transferred to ice. Cells were poured onto a filter paper, washed with 5 mL of mLV medium, and the filter paper with the cells was transferred to proliferation semi-solid mLV medium for 8 weeks (sample AR).

In the first experiment, global DNA methylation level was assessed in samples from lines 7P9a, 58P1a, and 58P5a, collected in triplicate before the experiment (T0) and during the cryopreservation process (T24, T48, T72, and after LN). In the second experiment, global DNA methylation level was assessed in samples from lines B5, B14, and B50, collected in triplicate before the experiment (T0), after 0.2 M maltose treatment (T48), and 8 weeks after recovery (AR), and compared to that of control cell masses that did not went through the cryopreservation process and were maintained by sequential subcultures (C samples).

### DNA Extraction

Genomic DNA was extracted using the modified CTAB method (Doyle and Doyle 1990). Calli (100-300 mg) were collected in an Eppendorf tube, frozen by immersion in LN and homogenized with a pestle by adding 600 μL of extraction buffer (2% CTAB, 20 mM EDTA, 1.4 M NaCl, 100 mM Tris-HCl pH 8.0, 1% PVP 4000 and 2% β-mercaptoethanol) preheated to 65°C. After incubating the tubes at 65°C for 30 min, 500 μL of isoamyl alcohol: chloroform (24: 1 v/v) were added, stirred gently for a few seconds and centrifuged at 6000 rpm for 10 min. The upper phase was transferred to tubes with 700 μL of isopropanol, mixed by inversion and after 1 h at −20 °C, centrifuged at 13000 rpm for 30 min at 4 °C. After removing the supernatant, the pellet was subjected to successive washes with 70% and 96% ethanol and allowed to dry. The pellet was then resuspended in 125 μL of 1X TE (10 mM Tris HCl pH 8.0 and 1 mM EDTA pH 8.0), and 37.5 μL of 5M NaCl, 50 μL of 5M KAc and 75 μL of 30% PEG were added, leaving precipitate the DNA for at least 2 h on ice. The tubes were subsequently centrifuged at 13000 rpm (15 min at 4 °C), the supernatant was removed, and the pellet dried in the speed vac for 5-10 min. Finally, DNA was resuspended in 30 μL of mQ water and incubated with 1 μL of RNase (10 mg/mL prepared from a stock of 100 mg/mL of RNase A from Qiagen, Darmstadt, Germany) at 37 °C for 1 h. Samples were stored at −20°C until use.

### Quantification of global DNA methylation

Global DNA methylation levels were quantified by using 100 ng of genomic DNA from each sample and MethylFlash Methylated DNA Quantification Kit (Colorimetric; Epigentek, NY). This ELISA-based procedure estimates 5-methyl-cytosine fraction in genomic DNA extracts by using specific antibodies. The colorimetric assay determines the amount of methylated DNA as it is proportional to the OD (optical density) intensity measured at 450 nm. After subtracting negative control readings from the readings for samples and standards, the percentage of methylated cytosines (%5-mC) in total DNA was calculated for each sample as a ratio of its OD relative to the standard OD. Data are presented as a mean ± SE.

### Determination of oxidative stress levels

Oxidative stress was studied in both T0 and T48 samples from 3 embryogenic lines (B5, B14 and B50). Superoxide radicals were determined by using 10 μM dihydroethidium (DHE) as described in Rodríguez-Serrano et al. (2012). As a negative control and to downplay background signals, samples were incubated for 1 h with 4 mM MnCl2 as a ROS scavenger, before adding the fluorochromes. DHE signal was captured as red fluorescence (490 nm excitation; 520 nm emission) observed in samples with a confocal laser scanning microscope (Fluoview FV10iW, OLYMPUS). For each line and time, 6 biological samples were analyzed, and the fluorescence intensity was quantified in all the cells from each sample using the ImageJ software. Values of fluorescence intensity/cell surface were then corrected by subtracting the average signal of samples incubated with the scavenger.

### Statistical analyses

Data from %5-mC and fluorescence intensity determinations were subjected to analysis of variance (ANOVA) using SPSS v25 (IBM Statistics, USA). When percentages data were not normal, data were subjected to angular transformation before analysis. For mean separation, Tukey-b tests were performed.

## Results

### Changes in global DNA methylation during the cryopreservation procedure

Analyses were performed in two independent experiments with 3 embryogenic lines each. In both experiments, we found that at T0 global DNA methylation was highly variable among the embryogenic lines tested.

In the first experiment, samples before (T0) and during the cryopreservation process (T24, T48, T72, and LN) were prepared from lines 7P9a, 58P1a and 58P5a. Initial (T0) global DNA methylation levels of the three embryogenic lines were significantly different (*p* = 0.027), since samples from the 58P5A genotype showed lower %5-mC values than those from the 7P1A genotype (Figure 2). However, on average we did not find significant differences among lines (*p* = 0.770), while global DNA methylation levels changed during the cryopreservation process (*p* = 0.002). Averaged global DNA methylation percentages decreased from 0.41 ± 0.13 to 0.28 ± 0.14 after culturing embryogenic lines in liquid media with 0.4 M maltose (T72), and to 0.19 ± 0.08 after immersion of samples in LN. This result was mainly explained by the trend observed in the embryogenic line 7P1A (Figure 2).

**Figure 2.**
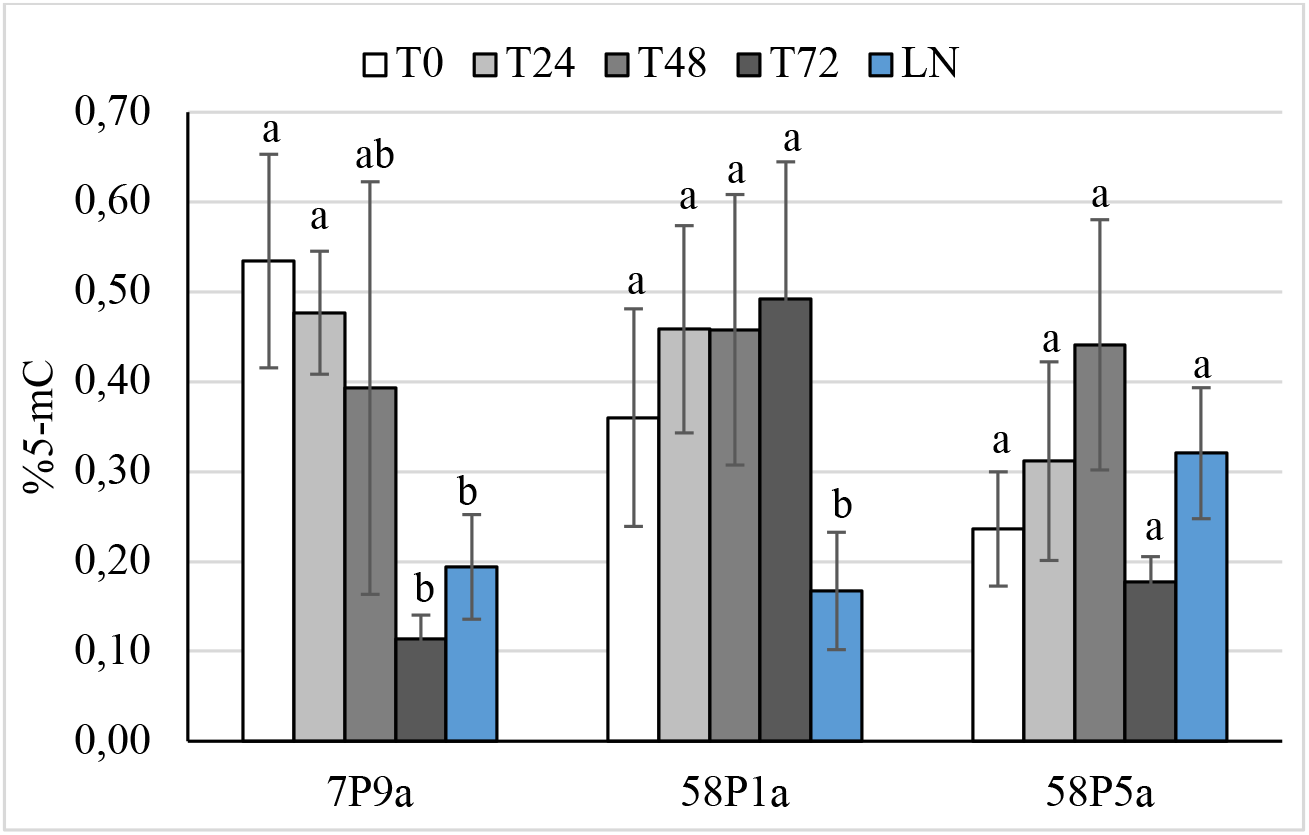
Global DNA methylation, assessed as percentage of methyl cytosine (%5-mC), in genomic DNA from three maritime pine embryogenic lines (7P9a, 58P1a, and 58P5a) sampled during the cryopreservation process. T0, before cryopreservation; T24, after cultured in liquid medium; T48, after culture in 0.2 M maltose; T72, after increasing maltose up to 0.4 M; LN, after being submerged in liquid nitrogen. Data are mean ± SE of at least 3 replicates; within each line, mean values followed of the same letter are not significantly different according to the Tukey-b test.

In a second experiment, samples at T0, T48, AR and C were prepared from B5, B14, and B50 embryogenic lines. Corroborating results from the first experiment, no significant differences were detected on average among lines (*p* = 0.288), but the ANOVA test estimated significant differences among samples throughout the cryopreservation process (*p* ≤ 0.001), and a significant interaction between the line and the time was also detected (*p* ≤ 0.001). Global DNA methylation decreased after a 24 h culture in 0.2 M maltose (T48 samples) for lines B5 and B14, while this pretreatment did not affect levels of % 5-mC in B50 embryogenic cells (Figure 3). After 8 weeks of culture, embryogenic cells of line B5 recovered from cryopreservation regained initial global DNA methylation percentages, while in B14 and B50 cell cultures methylation percentages did not increase after recovery (Figure 3). Not cryopreserved cell cultures of these two lines showed significantly higher global DNA methylation levels as compared to the initial values (Figure 3), which contrasted again with the significantly lower %5-mC values determined in the B5 cell line. It is noteworthy that, as observed in the first experiment, the three lines showed significantly different initial %5-mC values (*p* < 0.001), since we determined global DNA methylation levels of 0.53 ± 0.04, 0.69 ± 0.02, and 0.32 ± 0.01, for B5, B14, and B50, respectively.

**Figure 3.**
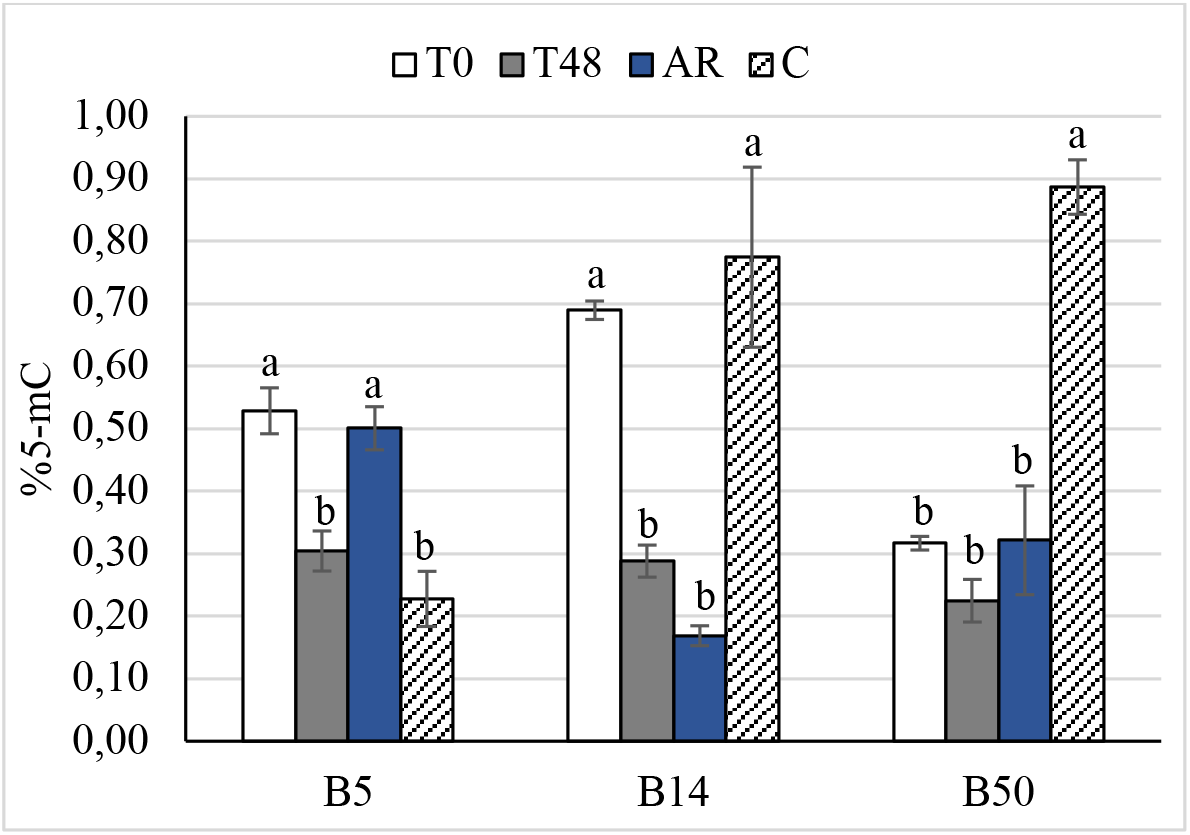
Global DNA methylation, assessed as percentage of methyl cytosine (%5-mC), in genomic DNA from maritime pine embryogenic lines B5, B14, and B50 sampled during the cryopreservation process. T0, before cryopreservation; T48, after culture in 0.2 M maltose; AR, after 8-week recovery; C, no cryopreserved. Data are mean ± SE of 3 samples. For each line, values followed of the same letter are not significantly different according to the Tukey test.

### Oxidative stress

The DHE fluorescence assay (Figure 4) to determine oxidative stress in the maritime pine embryogenic lines showed lower initial intensity/cell surface values in line B5 (*p* = 0.031) as compared to B14 and B50 lines (5.4 ± 3.2 front to 23.1 ± 4.4 and 22.8 ± 6.4, respectively). A 0.2 M maltose pretreatment slightly decreased ROS content of embryogenic cells in B14 and B50 lines (13.2 ± 6.1 and 5.6 ± 4.0), although these differences were not significant (*p* = 0.078). Fluorescence levels at T48 remained unaffected in B5 line (9.0 ± 2.7, *p* = 0.333).

**Figure 4.**
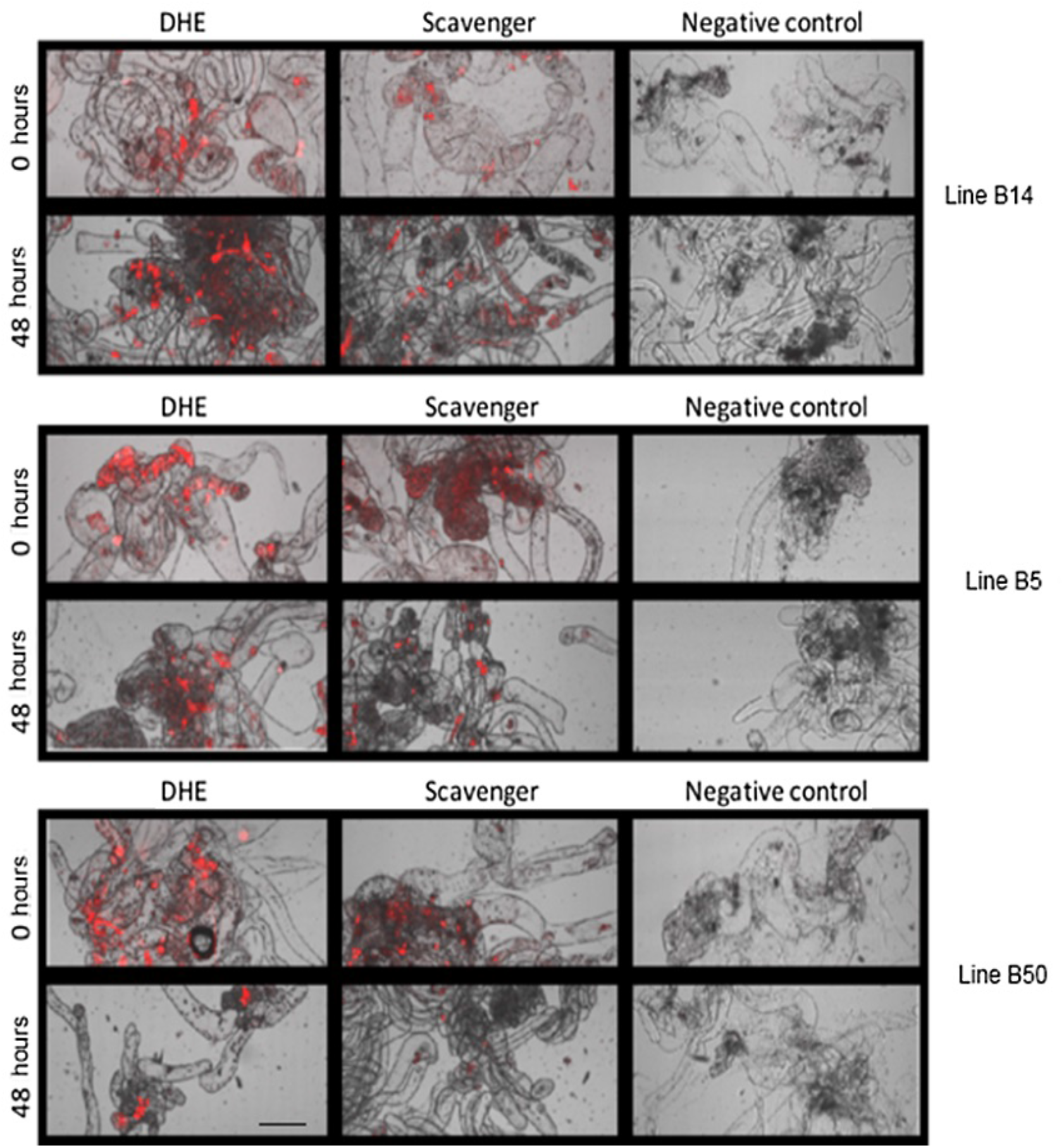
Examples of ROS quantification, as assessed by DHE fluorescence, in samples of maritime pine embryogenic lines B14, B5 and B50, before (T0) and after (T48) a maltose pretreatment in liquid media during the cryopreservation protocol. Basal levels were determined by incubating samples with 4 mM MnCl2 as ROS scavenger. Bar, 500 μm.

## Discussion

DNA-methylation levels varied among maritime pine embryogenic lines during the proliferation phase of the SE process (T0 samples). This fact might be due to the heterogeneous cell composition of the conifer embryogenic lines, as has been previously reported (Klimaszewska et al., 2016, Arrillaga et al. 2019). Nevertheless, a genotype influence should not be discarded, as has been described in scots pine (Alakärppä 2018). Note that the embryogenic lines employed in the experiments derived from different mother trees sampled in two populations of the species and lines from the same mother tree (58P1a and 58P5a) do not differ in their initial global DNA methylation.

In conifers, SE induction is usually accompanied by a drastic hypomethylation of DNA, enabling the expression of key genes involved in the initiation of the embryogenic process (De la Peña et al. 2015), whereas methylation levels increase during somatic embryo differentiation (Teyssier et al. 2013). Then, initial methylations rates observed in some of our embryogenic lines could also be associated with the specific developmental stage at which they were collected. A liquid culture pretreatment with high maltose concentration (T48 and T72) decreased global DNA methylation of maritime pine embryogenic calli. During this maltose pre-treatment (samples T48), ROS activity was not affected in maritime pine embryogenic cells, therefore ROS accumulation derived from increased osmotic stress during the cryopreservation process was not observed. Activation of transcriptional machinery of genes to alleviate osmotic stress might explain these results, since changes in methylation of genome areas containing genes involved in stress response, that were primarily hypomethylated, are well documented (Kovalchuk 2016). The antioxidant-related gene expression after cryopreservation was correlated to high survival rates in arabidopsis seedlings (Chen et al. 2015). Whether such changes at specific points may affect further expression of specific genes needs to be studied. As an example, in rice, osmotic stress caused DNA hypomethylation, and thereby expression activation, of genes involved in proline accumulation (Zhang et al. 2013). Further research is needed in order to learn if the specific demethylated genes remain the same, prior and after the cryopreservation process.

Lower levels of global DNA methylation were also detected after cryopreservation (LN samples). This agrees with previous reports in which demethylation events were the most frequent epigenetic changes observed when compared MSAP profiles of cryopreserved and non-cryopreserved control plant material (González-Benito et al. 2020). After thawing and transference to mLV medium, embryogenic lines of maritime pine proliferated without phenotypic or global DNA methylation status alterations, as compared to the same material at T0. Epigenetic changes induced during cryopreservation can be transient, as described by Zhang et al. (2020), who detected (by MSAP) 12.8% and 1.6% DNA methylation in kiwifruit cryo-derived shoots when cultured in vitro and in the cryo-derived plants after re-established in greenhouse conditions, respectively. Also, our results are in accordance with those obtained in mint (*Mentha x piperita* L) shoot apices cryopreservation, where significant epigenetic changes associated to several steps of the protocol were observed (Ibáñez et al. 2019). Global DNA methylation profiles of peach palm somatic embryogenic masses were influenced by both cryoprotectants and droplet-vitrification cryopreservation, but initial levels were recovered after regrowth (Heringer et al. 2013). Interestingly, cryopreserved seeds of maize showed some delay in germination, that was associated with changes in the proportion of DNA methylation in genes related to growth and development, being these changes depending on the germination stage and the cryopreservation treatment (Pérez et al. 2017). Hao et al. (2001) associated cryopreservation-induced demethylation events to the increased root capacity of apple shoots observed after cryopreservation. Furthermore, Nuc et al. (2016) studied DNA methylation after cryopreservation of embryonic axes isolated from seeds of pedunculate oak (*Quercus robur*) and European beech (*Fagus sylvatica*), and reported different methylation status depending on their tolerance (beech) or recalcitrance (oak) to dehydration. The loss of embryogenic potential observed in long-term subcultures of *Quercus alba* was associated by Corredoira et al. (2017) to an increase in DNA methylation, and therefore might account for the lower maturation rates observed in aged maritime pine embryogenic lines.

In conclusion, our results suggest that the maintenance of the embryogenic lines without cryopreservation but with periodical subcultures onto the same medium might increase methylation status of the DNA. In contrast, cryopreserved material did not show that increase, since although osmotic stress pretreatments and immersion in LN altered global DNA-methylation levels, maritime pine embryogenic cell masses recovered after thawing their initial methylation status.

## Acknowledgements

authors would like to thank to Álex Alborch for the technical assistance. This work was supported by the research project co-financed by MICINN and FEDER from the EU (AGL2013-47400-C4-04-R and AGL2016-76143-C4-01-R).

## Author’s contribution

Conceptualization IA; Material preparation, data collection and analysis IMP, MC, MTS, FD, PST, ES; Formal analysis and investigation ES, IA; Writing - original draft preparation IMP, JS, ES, IA; Writing - review and editing JS, ES, IA; Funding acquisition IA. All authors read and approved the final manuscript.

## References

Alakärppä E, Salo HM, Valledor L, Cañal MJ, Häggman H, Vuosku J (2018) Natural variation of DNA methylation and gene expression may determine local adaptations of Scots pine populations. J Exp Bot 69: 5293–5305 2018 doi:10.1093/jxb/ery292

Arrillaga I, Guevara MA, Muñoz-Bertomeu J, Lázaro-Gimeno D, Sáez-Laguna E, Díaz LM. Torralba L, Mendoza-Poudereux I, Segura J, Cervera MT (2014) Selection of haploid cell lines from megagametophyte cultures of maritime pine as a DNA source for massive sequencing of the species. Plant Cell Tiss Org Cult 118: 147–155. doi: 10/1007/s11240-014-0470-z

Arrillaga I, Morcillo M, Zanón I, Lario F, Segura J, Sales E (2019) New Approaches to Optimize Somatic Embryogenesis in Maritime Pine. Front. Plant Sci. 10:138. doi: 10.3389/fpls.2019.00138

Cano M, Morcillo M, Humánez A, Mendoza-Poudereux I, Alborch A, Segura J, Arrillaga I (2018) Maritime pine Pinus pinaster Aiton, In: S. M. Jain and P. Gupta (eds.), Step Wise Protocols for Somatic Embryogenesis of Important Woody Plants, Forestry Sciences 84, 167–180. Springer International Publishing AG. https://doi.org/10.1007/978-3-319-89483-6_13

Chen G-Q, Ren L, Zhang J, Reed BM, Zhang D, Shen X-h (2015) Cryopreservation affects ROS-induced oxidative stress and antioxidant response in Arabidopsis seedlings. Cryobiol 70 38–47

Corredoira E, Cano V. Bárány I, Solís MT, Rodríguez H, Vieitez AM, Risueño MC, Testillano MP (2017) Initiation of leaf somatic embryogenesis involves high pectin esterification, auxin accumulation and DNA demethylation in *Quercus alba*. J Plant Physiol. 213: 42–54. doi:10.1016/j.jplph.2017.02.012

De-la-Peña C, Nic-Can GI, Galaz-Ávalos RM, Avilez-Montalvo R, Loyola-Vargas VM (2015) The role of chromatin modifications in somatic embryogenesis in plants. Front Plant Sci 6: 635. doi:10.3389/fpls.2015.00635.

Doyle JJ, Doyle JL (1990) Isolation of plant DNA from fresh tissue. Focus 12: 13–15

Gallego-Bartolomé J (2020) DNA methylation in plants: mechanisms and tools for targeted manipulation. New Phytol 227: 38–44. doi: 10.1111/nph.16529

González-Benito ME, Ibáñez MA, Pirredda M, Mira S, Martín C (2020) Application of the MSAP technique to evaluate epigenetic changes in plant conservation. Int J Mol Sci 21: 7459. doi:10.3390/ijms21207459

Hao Y-J, Liu, Q-L, Deng X-X (2001) Effect of cryopreservation on apple genetic resources at morphological, chromosomal, and molecular levels. Cryobiol 43: 46–53

Heringer AS, Steinmacher DA, Fraga HPF, Vieira LN, Ree JF, Guerra MP (2013) Global DNA methylation profiles of somatic embryos of peach palm (*Bactris gasipaes* Kunth) are influenced by cryoprotectants and droplet-vitrification cryopreservation. Plant Cell Tiss Org Cult 114: 365–372

Huang H, Ullah F, Zhou D-X, Yi M, Zhao Y (2019) Mechanisms of ROS regulation of plant development and stress responses. Front Plant Sci 10: 800. https://www.frontiersin.org/article/10.3389/fpls.2019.00800

Humánez A, Blasco M, Brisa C, Segura J, Arrillaga I (2012) Somatic embryogenesis from different tissues of Spanish populations of maritime pine. Plant Cell Tiss Org Cult 111: 373–383. doi: 10.1007/s11240-012-0203-0

Ibáñez MA, Alvarez-Mari A, Rodríguez-Sanz H, Kremer C. González-Benito ME, Martín C (2019) Genetic and epigenetic stability of recovered mint apices after several steps of a cryopreservation protocol by encapsulation-dehydration. A new approach for epigenetic analysis. Plant Physiol Biochem 143: 299–307. doi.org/10.1016/j.plaphy.2019.08.026

Klimaszewska K, Hargreaves C, Lelu-Walter MA, Trontin JF (2016) Advances in conifer somatic embryogenesis since year 2000. In: In Vitro Embryogenesis in Higher Plants, Vol. 1359, eds M. A. Germanà and M. Lambardi (New York, NY: Humana Press), pp 131–162. doi: 10.1007/978-1-4939-3061-6-7

Kovalchuk I (2016) Transgenerational genome instability in Plants. In: Genome Stability, Elsevier Inc, pp 615–636. http://dx.doi.org/10.1016/B978-0-12-803309-8.00036-7

Kurdyukov S, Bullock M (2016) DNA methylation analysis: choosing the right method. Biology 5: 3. doi:10.3390/biology5010003

Lelu-Walter MA, Bernier-Cardou M, Klimaszewska K (2006) Simplified and improved somatic embryogenesis for clonal propagation of *Pinus pinaster* (Ait.). Plant Cell Rep 25: 767–776

Lelu-Walter MA, Klimaszewska K, Miguel C, Aronen T, Hargreaves C, Teyssier C (2016) Somatic embryogenesis for more effective breeding and deployment of improved varieties in *Pinus* spp.: bottlenecks and recent advances. In: Somatic Embryogenesis: Fundamental Aspects and Applications, eds V. M. Loyola-Vargas and N. Ochoa-Alejo (Cham: Springer International Publishing), pp 319–365. doi: 10.1007/978-3-319-33705-0_19

Liu X, Xu Y, Liu F, Pan Y, Miao L, Zhu Q, Tan S (2021) The feasibility of antioxidants avoiding oxidative damages from reactive oxygen species in cryopreservation. Front Chem 9: 648684. https://doi.org/10.3389/fchem.2021.648684

Marum L, Estêvão C, Oliveira M, Amâncio S, Rodrigues L, Miguel C (2004) Recovery of cryopreserved embryogenic cultures of maritime pine-effect of cryoprotectant and suspension density. CryoLett 25: 363–374

Marum L, Rocheta M, Maroco J, Oliveira M, Miguel C (2009) Analysis of genetic stability at SSR loci during somatic embryogenesis in maritime pine (*Pinus pinaster*). Plant Cell Rep 28: 673–682. https://doi.org/10.1007/s00299-008-0668-9

Niederhuth CE, Schmitz RJ (2017) Putting DNA methylation in context: from genomes to gene expression in plants. Biochem Biophys Acta - Gene Regulatory Mechanisms 1860: 149–156

Nuc K, Marszalek M, Pukacki PM (2016) Cryopreservation changes the DNA methylation of embryonic axes of *Quercus robur* and *Fagus sylvatica* seeds during in vitro culture. Trees 30: 1831–1841. doi 10.1007/s00468-016-1416-3

Osabe K, Clement JD, Bedon F, Pettolino FA, Ziolkowski L, Llewellyn J, Finnegan E, Wilson IW (2014) Genetic and DNA Methylation Changes in Cotton (*Gossypium*) Genotypes and Tissues. PLoS ONE 9(1): e86049. doi:10.1371/journal.pone.0086049

Park YS (2002) Implementation of conifer somatic embryogenesis in clonal forestry: technical requirements and deployment considerations. Ann For Sci 59: 651–656

Perez J, Araya-Valverde E, Garro G, Abdelnour-Esquivel A (2017) Analysis of Stress Indicators During Cryopreservation of Seeds of Landrace Maize (*Zea mays*). CryoLett 38: 445–454

Ren L, Zhang D, Chen G-Q, Reed BM, Shen X-H, & Chen H-Y (2015) Transcriptomic profiling revealed the regulatory mechanism of *Arabidopsis* seedlings response to oxidative stress from cryopreservation. Plant Cell Rep 34: 2161–2178. doi: 10.1007/s00299-015-1859-9

Rodríguez-Serrano M, Bárány I, Prem D, Coronado MJ, Risueño MC, Testillano PS (2012) NO, ROS, and cell death associated with caspase-like activity increase in stress-induced microspore embryogenesis of barley. J Exp Bot 63: 2007–2024

Teyssier C, Maury S, Beaufour M, Grondin C, Delaunay A, Le Metté C, Kevin A, Cadene M, Label P, Lelu-Walter MA (2014) In search of markers for somatic embryo maturation in hybrid larch (*Larix* × *eurolepis*): global DNA methylation and proteomic analyses. Physiol Plantarum 150: 271–291. doi:10.1111/ppl.12081

Tirnaz S, Batley J (2019) DNA Methylation: Toward Crop Disease Resistance Improvement. Trends Plant Sci 24:1137–1150. doi: 10.1016/j.tplants.2019.08.007

Zhang H, Lang Z, Zhu JK (2018) Dynamics and function of DNA methylation in plants. Nat Rev Mol Cell Biol 19: 489–506

Zhang CY, Wang NN, Zhang YH, Feng QZ, Yang CW, Liu B (2013) DNA methylation involved in proline accumulation in response to osmotic stress in rice (*Oryza sativa*). Genet Mol Res 12: 1269–1277. doi: 10.4238/2013

Zhang X-C, Bao W-W, Zhang A-L, Wang Q-C, Liu, Z-D (2020) Cryopreservation of shoot tips, evaluations of vegetative growth, and assessments of genetic and epigenetic changes in cryo-derived plants of *Actinidia* spp. Cryobiol 94: 18–25

